# Evidence for ancient selective sweeps followed by differentiation among three species of *Sphyrapicus* sapsuckers

**DOI:** 10.1101/2023.03.24.534187

**Authors:** Libby Natola, Darren Irwin

## Abstract

Speciation occurs when gene pools differentiate between populations, but that differentiation is often highly heterogeneous across the genome. Understanding what parts of the genome are more prone to differentiation can inform us about genomic regions and evolutionary processes that may be central to the speciation process. Here, we study genomic variation among three hybridizing species of North American woodpecker: red-breasted, red-naped, and yellow-bellied sapsuckers (*Sphyrapicus ruber, S. nuchalis,* and *S. varius*). We use whole genome resequencing to measure genetic variation among these species and to quantify how the level of differentiation varies across the genome. We find that regions of high relative differentiation between species (*F*_ST_) tend to have low absolute differentiation between species (π_B_), indicating that regions of high relative differentiation often have more recent between-population coalescence times than regions of low relative differentiation do. Most of the high-*F*_ST_ genomic windows are found on the Z chromosome, indicating this sex chromosome is particularly important in sapsucker differentiation and potentially speciation. These results are consistent with a model of speciation in which selective sweeps of globally advantageous variants spread among partly differentiated populations, followed by differential local adaptation of those same genomic regions. We propose that sapsucker speciation may have occurred primarily via this process occurring on the Z chromosomes, resulting in genetic incompatibilities involving divergent Z chromosomes.

## Introduction

The speciation process is responsible for the generation of all species diversity and is a major topic of study in evolutionary biology. Speciation occurs after some barrier reduces or precludes gene flow between populations, fracturing individuals into multiple isolated groups which then accumulate differences over time (Coyne & Orr, 2004; Price, 2008). Barriers may be caused by genetic incompatibilities (Huang et al., 2020), behavioral differences (Uy et al., 2018), ecological differences, or geographic barriers to dispersal (Nelson, 2006); and they could have existed in the past (J. T. Weir & Schluter, 2004) or be contemporary (Clark et al., 2010). Often, geographic separation acts as an initial cause of differentiation (Price 2008), allowing genetic and phenotypic differences to evolve that later cause reduced hybrid fitness if populations come into contact.

Many closely related groups in the intermediate stages of speciation today arose during the Pleistocene era, during which fluctuating periods of glaciation geographically isolated conspecific populations, initiating differentiation between groups that we describe today as species (Howell, 1952; Schluter & McPhail, 1992; J. T. Weir & Schluter, 2004). In North American bird populations, for example, a commonly occurring biogeographic pattern is the following: related groups (i.e., different species, subspecies, or clearly differentiated populations) are often geographically structured into a west coast group, an interior west group, and an eastern group, and these three types are often now in contact in western Canada (Weir and Schluter 2004). This pattern is a result of the Pleistocene glaciations separating forests into three major southern refugia, which have now expanded into western Canada.

An example of this biogeographic pattern is provided by three species of *Sphyrapicus* woodpeckers: red-breasted, red-naped, and yellow-bellied sapsuckers (*Sphyrapicus ruber, S. nuchalis,* and *S. varius*; Figure 1). Studies conducted using allozymes (Johnson & Zink 1983); mitochondrial DNA loci including Cytochrome-b (Cicero & Johnson 1995), COI (Natola & Burg 2018, Seneviratne et al. 2012), CR (Natola & Burg 2018); a Z-linked locus CHD1Z (Natola & Burg 2018, Seneviratne et al. 2012); and Genotyping-by-Sequencing (GBS) of the nuclear genome (Curtis 2017, Grossen et al. 2016, Natola et al. 2021, Seneviratne et al. 2016) all concur that the three forms are distinct based on genetic differentiation. Molecular dating based on mitochondrial DNA estimates that red-breasted and red-naped sapsuckers diverged about 0.32 million years ago (MYA), and their common ancestor diverged from yellow-bellied sapsuckers approximately 1.09 MYA (Weir & Schluter 2004), time frames over which there have been many cycles of glacial and interglacial periods (Menounos et al., 2009). Today, the species inhabit distinct habitats and climatic niches. All three species are associated with forested habitats containing quaking aspen groves (*Populus tremuloides*), but the yellow-bellied sapsuckers are found primarily in deciduous early-successional and riparian habitats, red-naped sapsuckers are most closely tied to higher elevation and more arid ponderosa pine (*Pinus ponderosa*) forests, and red-breasted sapsuckers inhabit coastal coniferous forests with high annual precipitation levels (Walters et al. 2020 a, b, c). There is a great deal of overlap in the areas in which these species can persist (Billerman et al. 2016, Natola & Burg 2018, Natola et al. 2023). These species hybridize where they come into contact, even forming a three-species hybrid zone (Natola et al. 2022) in central British Columbia, but the limited number of hybrids in that zone show that the three forms are stable species. Given the evidence of some current gene flow (Natola et al. 2022) as well as the deep genomic differentiation, it is a safe assumption that these three groups have experienced multiple cycles of geographic separation and hybridization during their path towards full species. Hence these sapsuckers provide an opportunity to examine the differentiation between three forms with a history of cycles of separation and contact.

**Figure 1.**
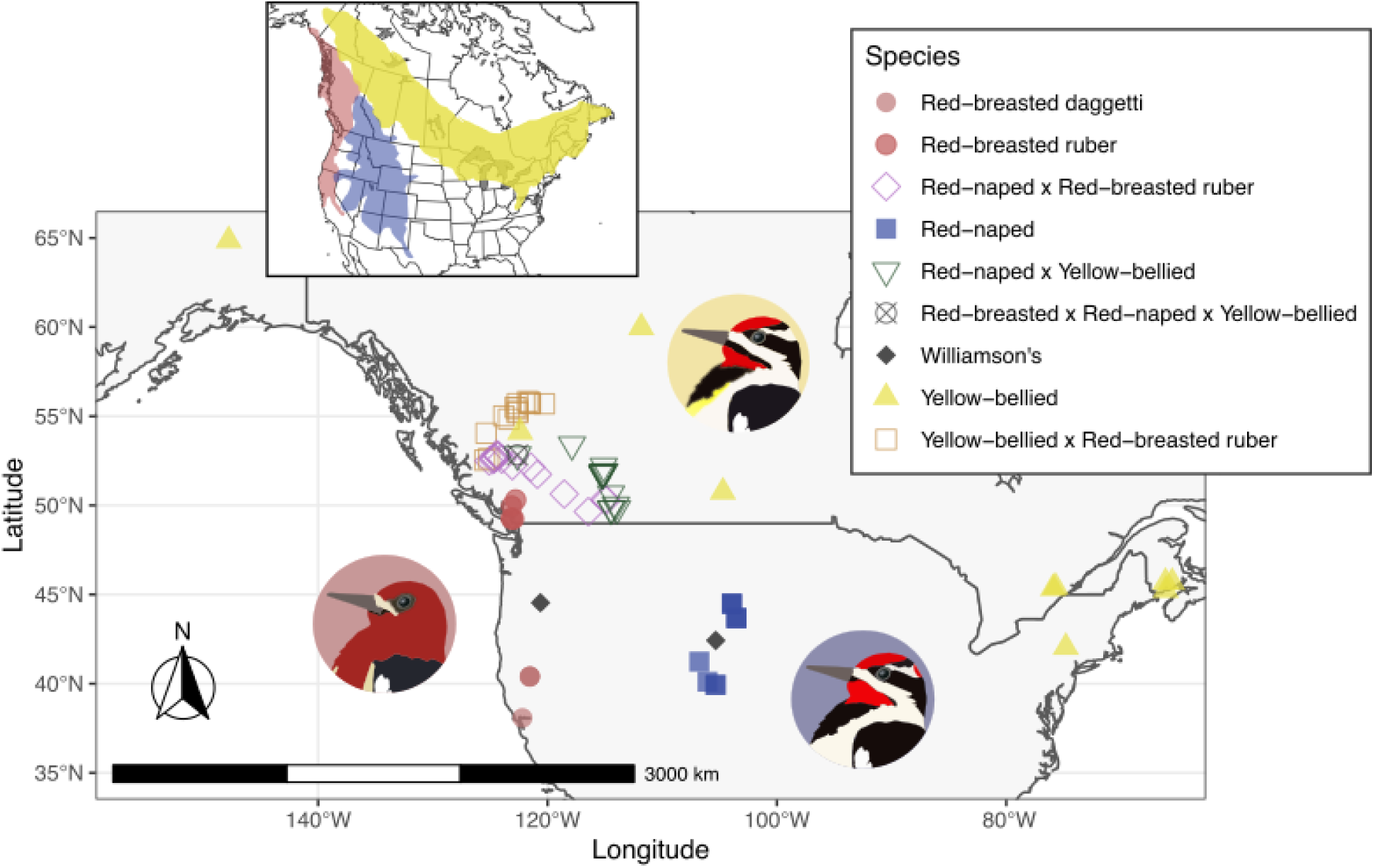
Sampling map of birds included in the WGS sequencing dataset (see Supplemental Table S1 for more details).

Genomic data are useful in speciation research because they reflect the evolutionary relationships between species, incorporating historical gene flow and population connectivity. The “genomic landscape of differentiation”, which describes the peaks and valleys of differentiation between species or populations along the genome, can help us identify regions that are most differentiated between species and might thereby have played a large role in speciation. There are different ways to measure differentiation, and comparing their genomic landscapes of variation can point to different evolutionary processes. By comparing relative allele frequency differentiation between populations (*F*_ST_), nucleotide distance between populations (π_B_; often referred to as “*D*_xy_”), and nucleotide diversity within populations (π_W_), we can test various differentiation models involving geographic separation, gene flow, and selection (Charlesworth, 1998; Cruickshank & Hahn, 2014; Irwin et al., 2016, 2018; T. L. Turner et al., 2005). There is a straightforward mathematical relationship between these aspects of diversity, with *F*_ST_ = (π_B_ -π_W_) / π_B_ when sample sizes are large and equal (Charlesworth 1998, Nei 1973). Holding π_W_ constant, higher π_B_ means higher *F*_ST_; holding π_B_ constant, higher π_W_ means lower *F*_ST_. Hence, we naively expect high-*F*_ST_ regions to be those with high π_B_ and low π_W_. To what degree this is true in real datasets depends on the details of the differentiation process.

Divergent selection acting upon a trait in different populations tends to cause greater allele frequency differentiation (i.e., *F*_ST_) in the loci which encode that trait compared to the rest of the genome. This process however is affected by the recombination landscape of the genome, such that a large genomic region of low recombination can be influenced by the combined effects of selection on multiple loci with the region. When genomic regions become differentiated through the combined effects of selection and low recombination, divergent loci might cause low fitness of hybrids, creating a barrier between the species. When populations are in contact, such divergent loci will tend to have less gene flow between populations than other parts of the genome do, resulting in comparatively high coalescence time—which is expected to be proportional to nucleotide distance (π_B_) when mutation rate is constant. This scenario leads to the expectation that regions of high *F*_ST_ will tend to also have high π_B_ (Cruickshank & Hahn, 2014; Han et al., 2017) (Figure 2A).

**Figure 2.**
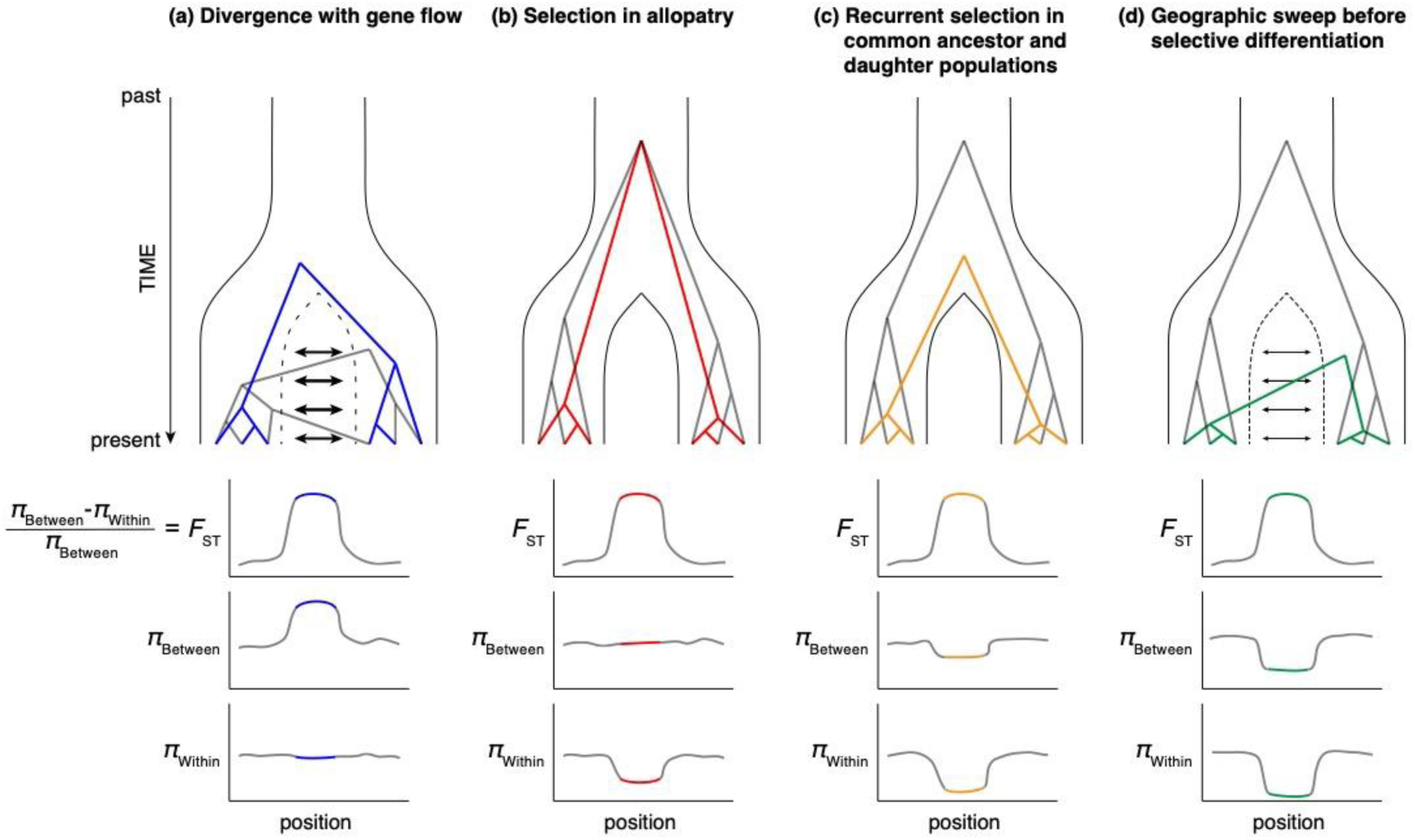
Illustration of four models of how selection can cause genomic regions to have high relative differentiation (*F*_ST_), along with the expected effects on between-group nucleotide distance (π_Between_) and within-group nucleotide diversity (π_Within_). Adapted from Irwin et al. (2018).

In contrast to this expectation, some pairs of differentiated populations do not show high π_B_ in the high *F*_ST_ regions of the genome (Cruickshank & Hahn 2014). In these cases, high *F*_ST_ is a result of low within-population nucleotide diversity (π_W_) rather than high between-population nucleotide distance. Proposed explanations for such a pattern center on selection (either positive or negative) in low-recombination regions of the genome resulting in especially low genetic diversity (Figure 2B). In some cases, there is even a negative relationship between *F*_ST_ and π_B_, a pattern for which two explanations have been proposed. One is that some regions of the genome have experienced selection-driven reductions in diversity in both the common ancestor and the two current populations (Cruickshank & Hahn 2014) (Figure 2C). Another is that adaptively advantageous mutations can sweep from one population to another across a hybrid zone, thereby lowering π_B_, and then subsequent local adaptation can increase *F*_ST_ (Delmore et al. 2015, Irwin et al. 2016) (Figure 2D). This latter model is called “sweep-before-differentiation” (Irwin et al. 2016, 2018). Differentiated yet hybridizing taxa such as the three species of *Sphyrapicus* sapsuckers provide excellent opportunities to examine the potential role of past gene flow in shaping the currently observed genomic landscapes of differentiation.

In addition to the influence of past gene flow and selection, genomic landscapes of differentiation are influenced by structural aspects of the genome (Noor et al., 2001; Payseur & Rieseberg, 2016; Rieseberg, 2001; Sætre et al., 1997), including sex chromosomes and inversions. In ZW sex chromosome systems such as birds (where females are the heterogametic sex), the Z chromosome often shows higher *F*_ST_ than the autosomes (Irwin, 2018; Qvarnström & Bailey, 2009). This is likely due to the smaller effective population size and reduced recombination, supplemented by greater potential for selective differentiation. Chromosomal inversions can also accumulate differentiation more rapidly than the rest of the genome, and they frequently act as isolating barriers (Hooper et al., 2018; Kirkpatrick & Barrett, 2015): crossing over in the inverted region of individuals with hybrid karyotypes may cause aneuploid gametes, reducing hybrid fitness (Hooper & Price, 2017; Kirkpatrick & Barton, 2006). Lack of recombination makes inversions a likely refuge for alleles that contribute to local adaptation (Coyne and Orr 2004), as in dune and non-dune prairie sunflowers (*Helianthus petiolaris*) (Huang et al., 2020; Todesco et al., 2020).

Here, we use whole genome resequencing (WGS) to measure and examine the genomic landscapes of differentiation between the three hybridizing *Sphyrapicus* sapsucker species. We use these results to determine which parts of the genome have high differentiation and whether these regions tend to have higher, similar, or lower nucleotide distances compared to the rest of the genome. We also examine genotypes of hybrids to infer whether there is evidence of recombination in highly differentiated regions, which would suggest that they are not inversions and that hybrids can reproduce successfully. The results from this analysis provide insight into the role of gene flow and selection during the speciation process in this three-species group that is biogeographically representative of many other taxa in North America.

## Materials and Methods

### Whole Genome Sequencing

#### Sample selection

Seventy-eight sapsucker samples were included in our WGS analysis: 10 allopatric individuals from each species, 15 hybrid zone individuals from each species cross, one putative 3-species hybrid, and two Williamson’s Sapsuckers (*Sphyrapicus thyroideus*) included as outgroups (Figure 1, Supplemental Table S1). We selected hybrid zone individuals to represent a continuum of admixture proportions between each species pair based on ancestry coefficient values from prior GBS research (Natola et al., 2021, 2022; Seneviratne et al., 2016). Samples were tissues from vouchered museum specimens and blood from wild-caught birds collected by Sampath Seneviratne (Seneviratne et al., 2012, 2016) and included similar proportions of both male and female birds (Supplemental Table S1).

#### DNA extraction, library preparation

We extracted DNA from all samples using a standard phenol-chloroform extraction procedure. We then quantified DNA concentrations using a Qubit fluorometer and submitted 100 ng DNA to Genome Quebec where technicians prepared shotgun DNA libraries using the NEB Ultra II (New England Biolabs) library preparation kit and then sequenced the libraries using four Illumina NovaSeq6000 S4 PE150 lanes.

#### Read processing

Sequencing resulted in 2.73 billion reads, with an average Phred quality score of 35. We used Trimmomatic v 0.38 (Bolger et al., 2014) to trim reads, and we then used BWA v 0.7.17 (Li, 2013) to align reads to our previously published red-breasted sapsucker reference genome (Natola et al. 2022). This reference genome is that of a female red-breasted sapsucker collected in Juneau, Alaska, and was assembled using long-read sequences and high sequencing depth (172x), and contigs were assembled into scaffolds based on a golden-fronted Woodpecker (*Melanerpes aurifrons*) reference genome (Wiley & Miller, 2020). After aligning reads, we added read group headers, converted to bam format, sorted, and indexed our reads using Picard v 1.97 (http://broadinstitute.github.io/picard/). Using samtools v 1.12 (Li et al., 2009), we merged the paired-end and un-paired bam files and indexed. We used the bamqc function from Qualimap v 2.2.2a (Okonechnikov et al., 2016) to assess quality of alignments. We created a corrected coverage measure from the bamqc files by dividing each scaffold’s read coverage per individual by that individual’s overall mean coverage. With this, we were able to identify scaffolds likely to be on the Z chromosome if known males had approximately twice the normalized coverage (normalized coverage = 1) as females (normalized coverage = 0.5), or on the W chromosome if known males had approximately zero normalized coverage compared to females’ 0.5 X coverage. We renumbered all Z scaffolds as their original golden-fronted woodpecker mapped scaffold number + 1000 and all W scaffolds as their original scaffold + 2000 (i.e., all scaffolds on the Z are labelled as between 1000 and 1999 and all W scaffolds are between 2000 and 2999). We then marked PCR (Polymerase Chain Reaction) duplicates in Picard 1.97 before identifying problematic regions, realigning, and fixing mate-pair coordinates using GATK v 3.8 (Mckenna et al., 2010) with scripts adapted from Aguillon et al. (2021). Finally, SNPs (Single Nucleotide Polymorphisms) were filtered in GATK by removing those with QD < 2.0, FS > 40.0, MQ < 20.0, and haplotype score > 12.0 settings, then in VCFtools v 0.1.16 (Danecek et al., 2011) with the < 20%, minor allele frequency > 5%, 3 < depth ≤ 50, and biallelic filters. We also created a dataset with all variant and invariant sites sequenced in the WGS dataset per scaffold, using the GATK v 4.0 (Mckenna et al., 2010) HaplotypeCaller tool. Then, we split the data into individual scaffolds and filtered with the same GATK 3.8 filtration protocol as the previous dataset, and similar VCFtools filtration with the exceptions of no minor allele frequency or minimum alleles filters to preserve invariant sites. Finally, we used the GATK 3.8 GenotypeGVCFs tool to include all variant and invariant sites across all individuals into one VCF file for each scaffold.

### Analyses

#### Species differentiation

To quantify mean site-wise genomic differentiation among the three species we calculated Weir & Cockerham’s (1984) pairwise weighted average *F*_ST_ in VCFtools using only allopatric samples, as we know differentiation is lower in hybrid zones and not representative of birds across the majority of their distribution. To visualize clusters of genomic similarity among the sapsuckers we constructed a PCA (Principal Component Analyses) of variation among individual genotypes using the SNPRelate package (Zheng et al., 2012) in the R programming language (R Core Team 2021). We calculated pairwise *F*_ST_ for all SNPs in 50 kb windows using VCFtools, then plotted the windowed average *F*_ST_ values using the package qqman (S. D. Turner, 2018) in R. We plotted *F*_ST_ (relative differentiation between populations), π_B_ (absolute differentiation between populations), and π_W_ (within population nucleotide diversity) across 50 kb non-overlapping sliding windows using allopatric individuals on all autosome and Z scaffolds > 50 kb; these together are referred to hereafter as the “genomic landscape dataset”. The total number of 50 kb windows included in these analyses was 18,550. We created these with scripts from Irwin et al. (2018) adapted in R and the Julia programming language (Bezanson et al. 2017). We separated autosomal and Z chromosome scaffolds and calculated *F*_ST_ using VCFtools. We calculated the differences between *F*_ST_ values on the autosomes versus Z chromosomes for each species pair using Welch’s two-sample t-tests and plotted the 50kb window means for *F*_ST_ vs. π_B_ for each species pair, while indicating the differences between differentiation on autosomes vs. Z chromosomes. We calculated Spearman’s rank correlation between *F*_ST_ and π_B_ for autosomes and Z chromosomes separately in R. We calculated the adjusted Z diversity, π_W,Z_*, as π_W_ /1.1 to account for the higher mutation rate on the Z chromosome (Irwin 2018), and then took the ratio of π_W,Z_*/π_W,A_ (π_W_ on the autosomes) to determine if diversity is comparable between Z chromosomes and autosomes. Mathematically, π_W,Z_*/π_W,A_ values < 0.5625 cannot be explained by even the most extreme asymmetries in variance of male and female reproductive success (e.g., harem or lek mating structures) (Charlesworth, 2001; Irwin, 2018).

#### Genotype × individual plots

To visualize Z-chromosome genotypes across many variant sites and individuals we created genotype-by-individual plots using the fixed (species-diagnostic, *F*_ST_ = 1) SNPs from each species pair per scaffold, and plotted these using Julia scripts that call the GenomicDiversity.jl package (Irwin et al. 2025). We removed all females to avoid complexity created by hemizygosity. We note that with this method SNPs are included in the three-way comparison that are fixed for differences between only two of the three species. To make plots easier to read, we thinned SNPs to only show SNPs with no missing genotypes and that are 100kb from other included SNPs.

## Results

Our data present clear evidence for strong differentiation between the three species. In the PCA (Figure 3), the major axis of variation (PC1) accounts for 14.77% of the variation and separates yellow-bellied sapsuckers from the other two species. The second axis of variation (PC2) accounts for 3.17% of the variation and separates red-breasted and red-naped sapsuckers. The weighted average pairwise *F*_ST_ values calculated for each species pair reflect these relationships: all three species pairs have clear differentiation (Table 1), and *F*_ST_ is highest in comparisons involving yellow-bellied sapsuckers.

**Figure 3.**
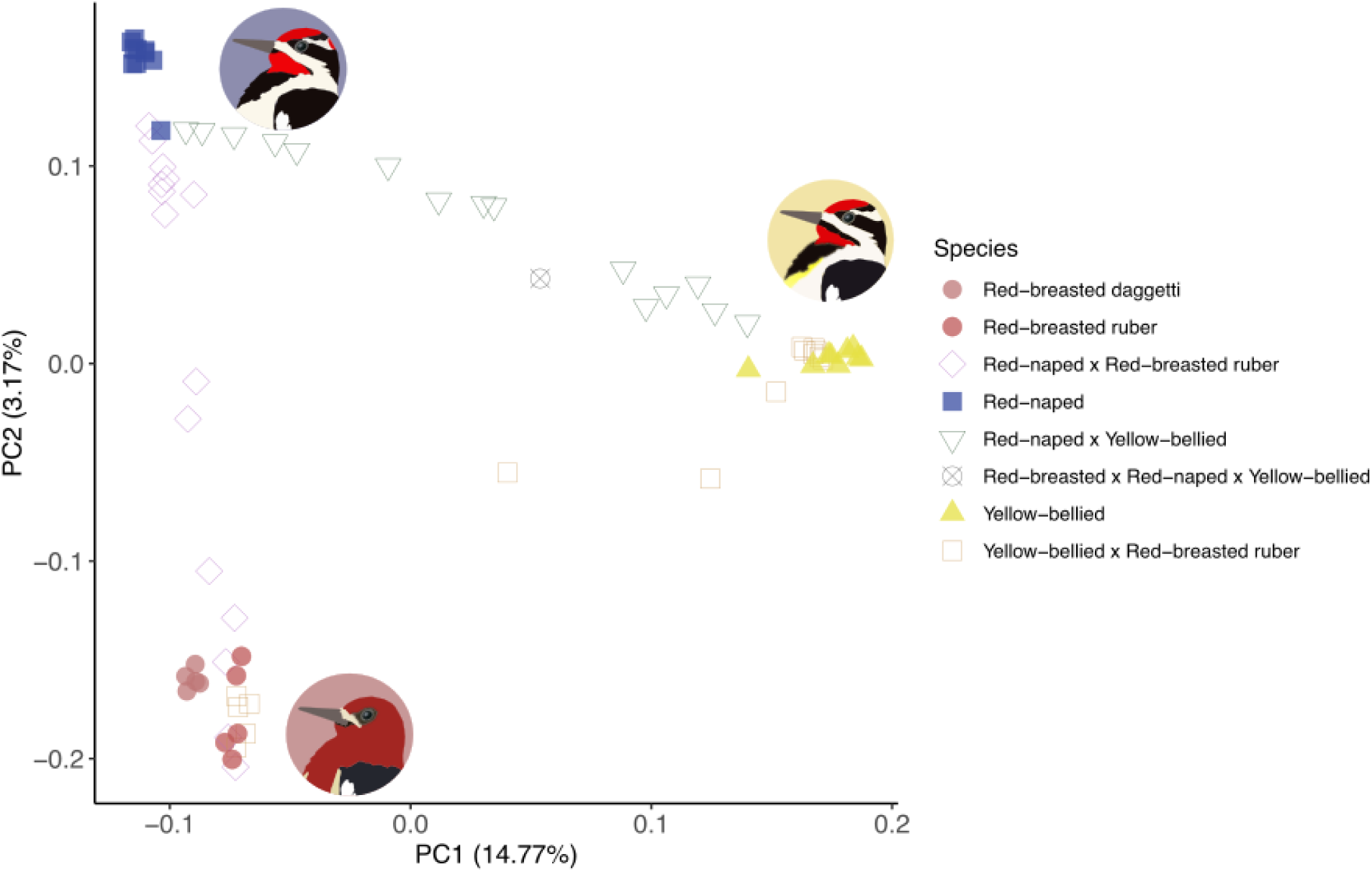
PCA of WGS genotypic data. Yellow dots indicate allopatric yellow-bellied sapsuckers, red dots indicate allopatric red-breasted, blue dots are allopatric red-naped. Orange dots show birds sampled in yellow-bellied × red-breasted sympatry, purple denote red-breasted × red-naped hybrid zone birds, and green dots show birds sampled in red-naped × yellow-bellied sympatry.

**Table 1.** Comparisons between autosomes (A) and Z chromosomes (Z) of the measurements *F*_ST_ and π_B_ between species (a), and π_W_ within species (b). We include adjusted Z diversity (π_W,Z_*) to correct for higher mutation rate on Z, and compare nucleotide differentiation on the Z vs autosome. Measurements were calculated as means of values from 50kb sliding windows from the genomic landscape dataset.

| Species pair | $F_{ST,A}$ | $F_{ST,Z}$ | $p$ value | $\pi_{B,A}$ | $\pi_{B,Z}$ | $p$ value | $\pi_{B,Z}^*$ |
| --- | --- | --- | --- | --- | --- | --- | --- |
| Red-naped/red-breasted | 0.098 | 0.412 | $p < 0.0001$ | 0.00321 | 0.00213 | $p < 0.0001$ | 0.00194 |
| Yellow-bellied/red-breasted | 0.209 | 0.546 | $p < 0.0001$ | 0.00478 | 0.00361 | $p < 0.0001$ | 0.00328 |
| Yellow-bellied/red-naped | 0.267 | 0.573 | $p < 0.0001$ | 0.00483 | 0.00359 | $p < 0.0001$ | 0.00326 |

b.
| Species | $\pi_{W,A}$ | $\pi_{W,Z}$ | $p$ value | $\pi_{W,Z}^*$ | $\pi_{W,Z}^*/\pi_{W,A}$ |
| --- | --- | --- | --- | --- | --- |
| Red-naped | 0.00270 | 0.00136 | $p < 0.0001$ | 0.00124 | 0.458 |
| Red-breasted | 0.00316 | 0.00172 | $p < 0.0001$ | 0.00156 | 0.496 |
| Yellow-bellied | 0.00458 | 0.00258 | $p < 0.0001$ | 0.00235 | 0.512 |

The windowed *F*_ST_ plots show many windows with low *F*_ST_ values (< 0.1) between red-breasted and red-naped sapsuckers, and overall higher average *F*_ST_ values in the species pairs involving yellow-bellied sapsuckers (Figure 4). The scaffolds with the highest average pairwise *F*_ST_ values occur in large peaks that are largely concurrent in each species pair (Figure 4, Supplemental Figures S1, S2). The most obvious pattern in these plots is the extremely high *F*_ST_ (many windowed values close to 1) across most of the Z chromosome. The weighted Weir and Cockerham *F*_ST_ of all SNPs (9,205,747 sites) on autosomes is 0.28 between red-naped and yellow-bellied, 0.22 between red-breasted and yellow-bellied, and 0.08 between red-breasted and red-naped. In contrast, on the Z scaffolds (655,683 sites) those measures are 0.47, 0.42, and 0.26, respectively. Whereas the autosomes contain 9,205,747 SNPs over 1.14 Gbp, only 8 of these are fixed between red-breasted and red-naped sapsuckers (0.000087 % of all variant sites), 26,347 (0.29%) are fixed between red-naped and yellow-bellied, and 20,361 (0.22%) are fixed between red-breasted and yellow-bellied. In contrast, on the Z chromosome scaffolds, which contain 655,683 polymorphic sites over 127 Mbp, 16,097 (2.5% of all variant sites), 66,906 (10.2%), and 64,873 (9.9%) are fixed between our samples of red-breasted/red-naped, red-naped/yellow-bellied, and red-breasted/yellow-bellied sapsuckers, respectively.

**Figure 4.**
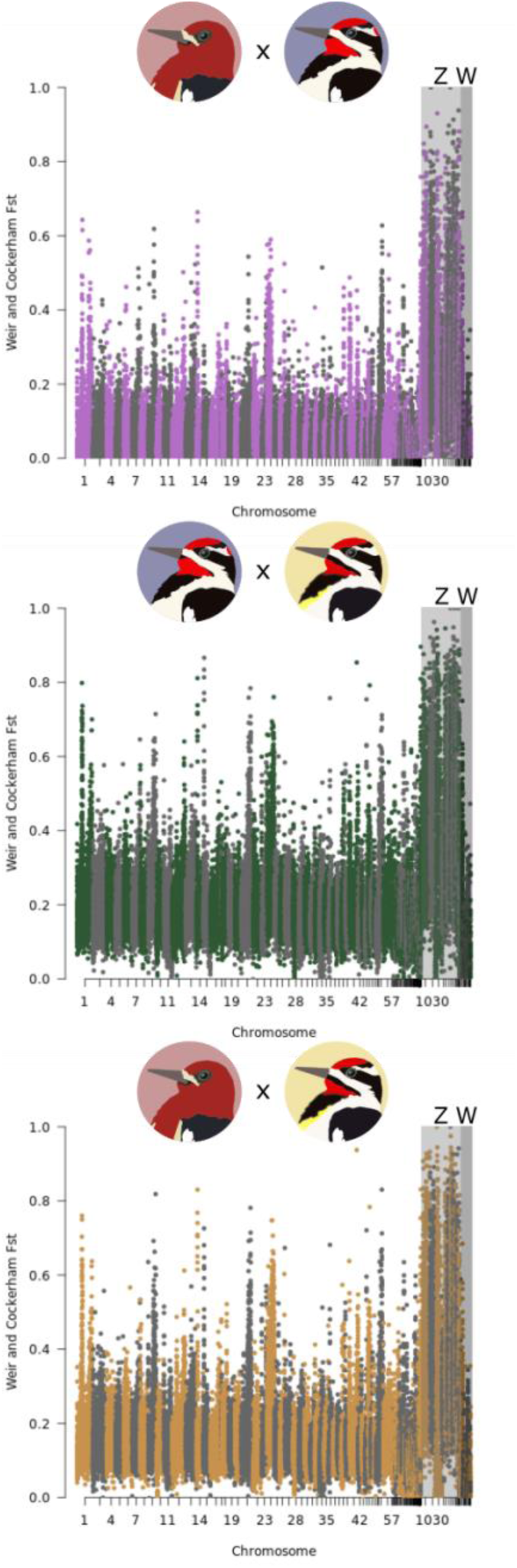
Genomic variation in pairwise *F*_ST_ between (a) red-breasted and red-naped sapsuckers, (b) red-naped and yellow-bellied sapsuckers, and (c) red-breasted and yellow-bellied sapsuckers. Each dot represents a weighted average over a 25 kb window. Light grey box shading indicates scaffolds identified as likely to be on the Z chromosome and darker grey box shading designates those on the W chromosome.

The mathematical relationship between *F*_ST_ and π_W_ (see Introduction) leads to the expectation that *F*_ST_ will tend to increase as π_W_ decreases, and that with π_W_ held constant regions of high *F*_ST_ will have high π_B_. The sapsucker comparisons show the opposite relationship, such that genomic regions of high *F*_ST_ tend to have low π_B_ with Spearman’s correlations indicating strong negative relationships between *F*_ST_ and πB in all comparisons of species, autosomes, and Z chromosomes (−0.264 ≤ ρ ≤ −0.544, all *p*-values < 0.0001; Figure 5). Low πB in high *F*_ST_ regions can occur if π_W_ is proportionally lower than π_B_. Biologically, this can be explained by selective sweeps in a common ancestor (lowering π_B_) and subsequent sweeps in the daughter species (lowering π_W_ and driving up *F*ST) (Cruickshank & Hahn, 2014) (Figure 2C), or a selective sweep across hybridizing populations (lowering π_B_) followed by local differentiation at these loci between populations (causing extremely low π_W_, therefore increasing *F*_ST_) (Irwin et al., 2016, 2018) (Figure 2D).

**Figure 5.**
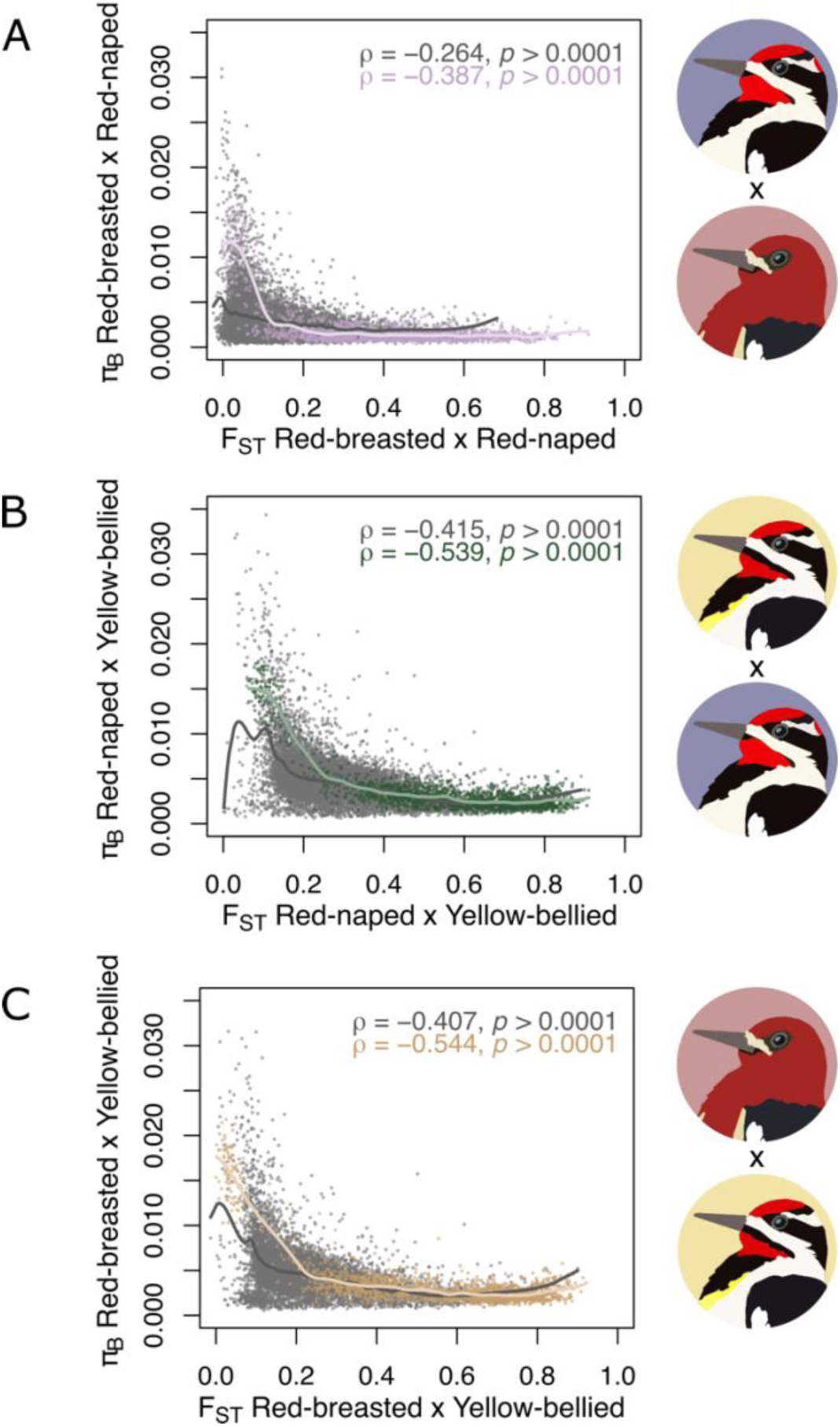
Comparisons of mean *F*_ST_ and π_B_ measured in 50 kb sliding windows for red-breasted and red-naped (A), red-naped and yellow-bellied (B), and red-breasted and yellow-bellied (C). Autosomal loci are shown in grey, and scaffolds presumed to be on the Z chromosome are shown in color. Spearman’s correlation rho and p values are shown for each species pair and chromosome category, with autosomal loci values written in grey text and Z chromosomal loci in color.

Windowed *F*_ST_ values among sapsucker species are much higher on the Z chromosome scaffolds than on the autosomes, within-group variation (π_W_) is lower, and between-group nucleotide distance (π_B_) is lower on the Z chromosome (Welch’s two-sample t-tests of the genomic landscape dataset including autosome and Z chromosome scaffolds > 50 kb: all *p*-values < 0.0001; Table 1, Figure 5). These data underscore the role of recurrent selection upon the Z chromosome. Indeed, in all three species, the ratio of adjusted π_W,Z_ to π_W,A_ is below the theoretical minimum of 0.5625 that can be explained without invoking selection (Charlesworth, 2001; Irwin, 2018). Within species, π_W_ is consistently significantly lower on the Z chromosome than on autosomes (Table 1). This is shown in the neighbor-joining trees of phylogenetic relationships between sapsuckers in Figure 6, which show shallower branching and proportionally shorter grey boxes on the Z than the autosomes. This indicates the high *F_ST_* regions have more recent coalescence times than the low-*F_ST_* regions.

**Figure 6.**
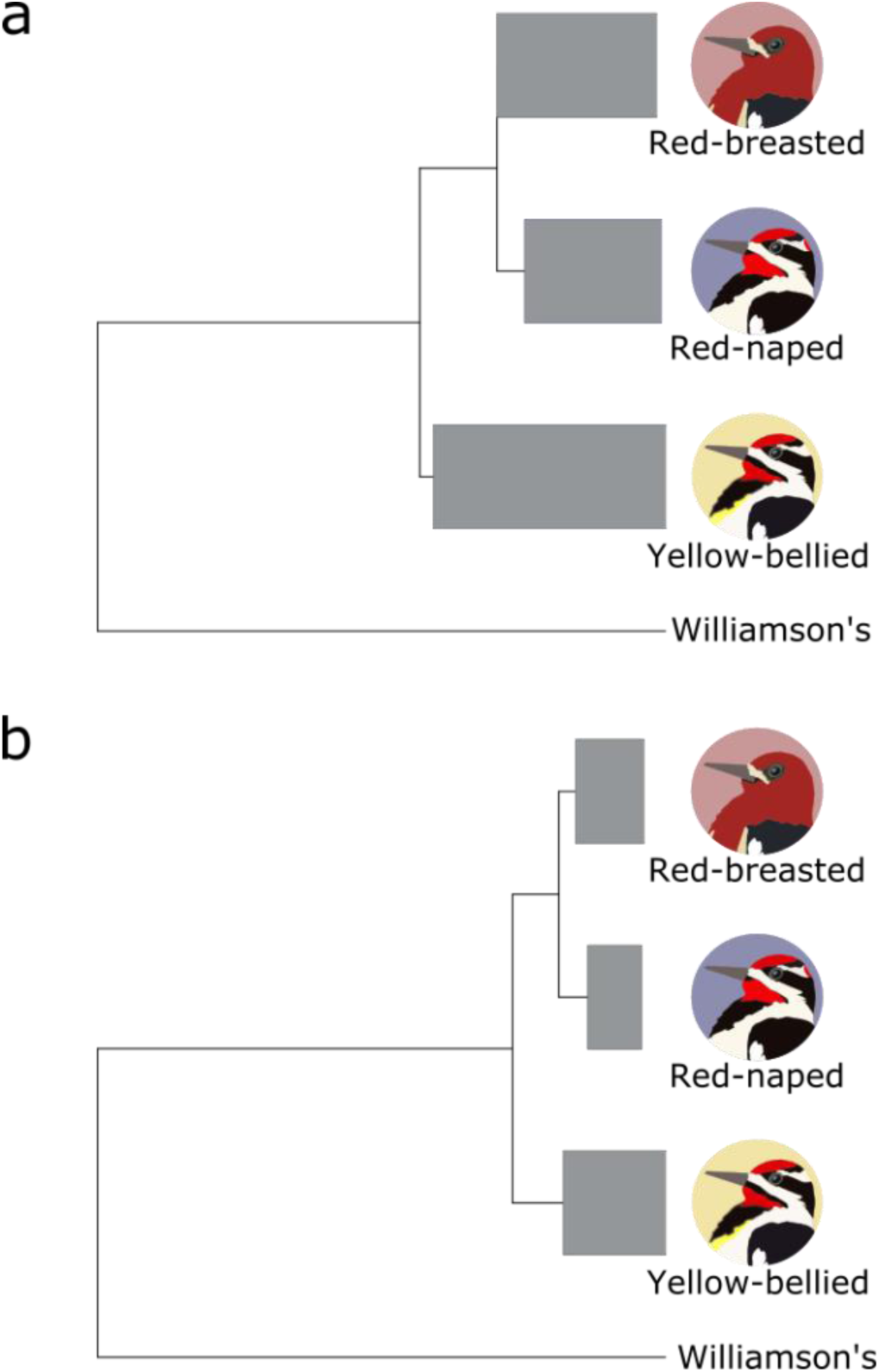
Neighbor-joining trees illustrating phylogenetic relationships among sapsuckers for autosomal (a), and Z-linked regions (b). Tree branch lengths are based on average between-taxon nucleotide distances (π_B_) and within-taxon between-individual nucleotide distances (π_W_, displayed as length of grey boxes).

As we have seen in Figures 3 and 4, there are strong differences in allele frequencies between the species, particular in the Z chromosome where windowed *F*_ST_ is often close to 1. In Figure 7 we use genotype-by-individual plots of two Z-chromosome scaffolds (together accounting for more than 47.0 Mbp) to illustrate the clear differentiation of these regions and to examine whether there is evidence of recombination in hybrids. The SNPs shown for each species pair have been filtered for those that are fixed between the species, for which there are no missing genotypes, and that are separated by at least 100,000 bp of chromosomal distance. Despite these strong filters, there are abundant SNPs that have fixed differences for each species pair. This allows a close look for evidence of recombination in hybrids, for which there is much evidence—this can be seen in hybrid individuals which have some long segments of heterozygosity and then other long segments of homozygosity. These can be explained as the result of past crossing-over and recombination during meiosis in individuals that have two Z chromosomes (i.e., generally males in birds) of different types.

**Figure 7.**
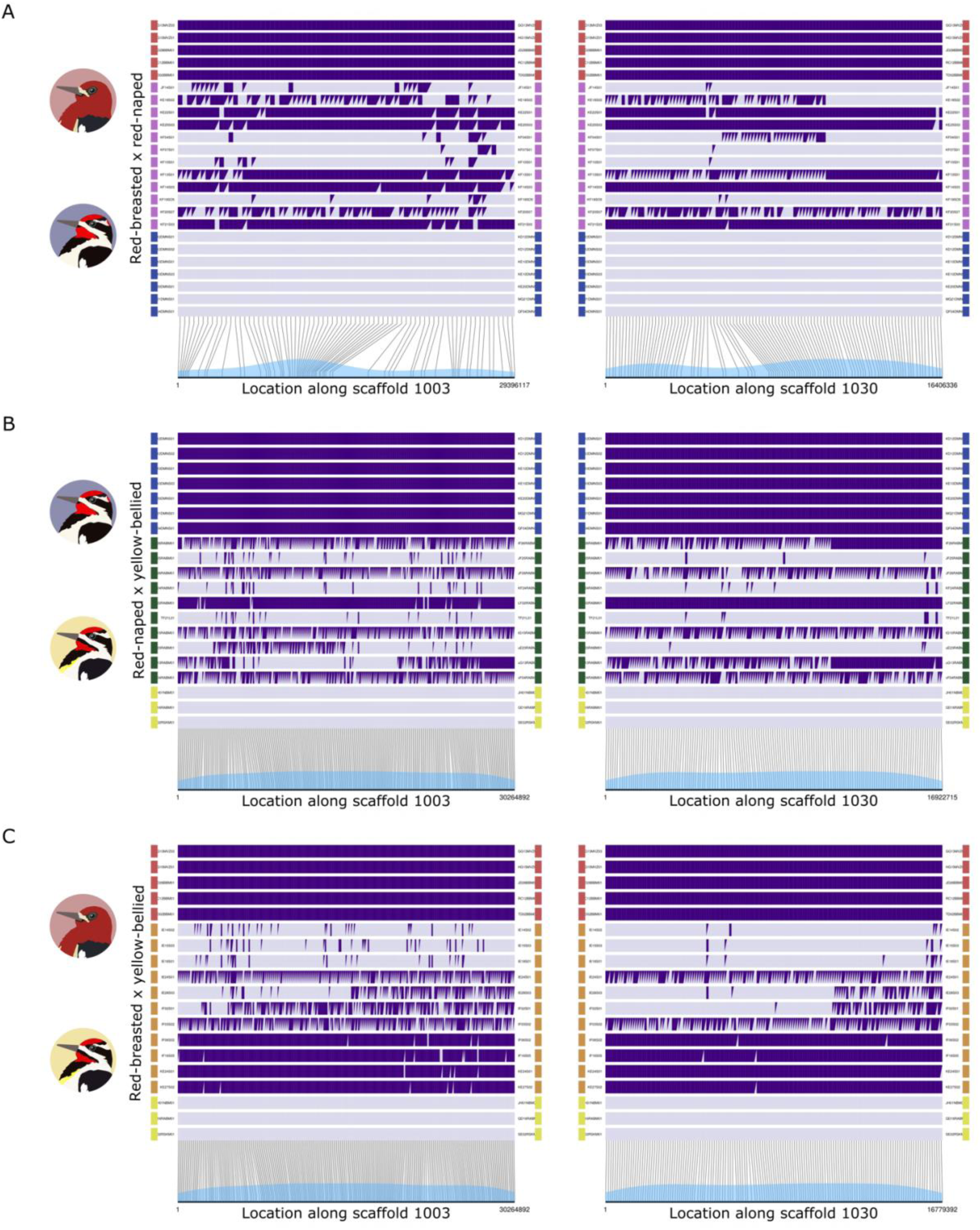
Genotype by individual plots showing strong Z chromosome differentiation between pairs of *Sphyrapicus* sapsucker species and clear evidence of recombination in hybrids. Panel A shows red-breasted / red-naped sapsuckers, panel B shows red-naped / yellow-bellied sapsuckers, and panel C shows red-breasted / red-naped sapsuckers. Two Z-chromosome scaffolds are shown: scaffold 1003 on left and scaffold 1030 on right. Rows within each panel correspond to individuals, with colors on left and right indicating the species or hybrid group they belong to: red-breasted (red), red-breasted × red-naped hybrid zone (purple), red-naped (blue), red-naped × yellow-bellied hybrid zone (green), red-breasted × yellow-bellied hybrid zone (orange), or yellow-bellied sapsuckers (yellow). Each genotype is depicted as either homozygous (purple rectangle or light purple rectangle) or heterozygous (rectangle split diagonally into a purple and light purple triangle). Alleles colored dark or light purple can differ between panels A, B, and C, with dark purple assigned to the first group in each comparison. Lines on the bottom of the plot indicate chromosomal position of each genotype along the scaffold. Only males are included on these plots, since females are hemizygous for the Z chromosome.

## Discussion

### Sapsucker relationships

We find overall patterns of autosomal differentiation among sapsuckers characterized by generally low *F*_ST_, punctuated by regions of high *F*_ST_ with low π_W_ and π_B_. These patterns are similar to several other sister taxa of North American boreal bird species, including mourning and MacGillivray’s warblers (*Geothlypis philadelphia, G. tolmiei*), myrtle and Audubon’s yellow-rumped warblers (*Setophaga coronata coronata, S. c. auduboni*), and Townsend’s, hooded, and black-throated green warblers (*Setophaga townsendi*, *S. cintra*, *S. virens*), which all show low π_B_ in high *F_ST_* regions (Irwin et al., 2018; Wang et al., 2021). The sapsucker genome scans show this pattern to an even stronger degree. Our results agree with existing literature in describing red-breasted and red-naped sapsuckers as sister species, with yellow-bellied sapsuckers positioned as an outgroup to this species pair (Cicero & Johnson, 1995; Grossen et al., 2020; Johnson & Zink, 1983; L. Natola & Burg, 2018). While many loci show remarkably low differentiation between red-breasted and red-naped sapsuckers (Cicero & Johnson, 1995; L. Natola & Burg, 2018), our genome-wide data show that the two exhibit high differentiation across large genomic distances on the Z chromosome. Our genome-wide mean pairwise *F*_ST_ calculations indicate higher differentiation between red-naped and yellow-bellied sapsuckers than between red-breasted and yellow-bellied. Lower within-population genetic diversity can lead to the higher relative between-population diversity we observe in the *F*_ST_ values. This might be a result of the closer geographic distances between red-naped sapsuckers used in this study, or of a species-wide pattern of low within-species genetic diversity.

### The genomic differentiation landscape and Sphyrapicus evolutionary history

One might expect because red-breasted and red-naped sapsuckers are more closely related their genomes would be similar (have low *F*_ST_ values) in the regions where each and the yellow-bellied lineages diverged, but regions of high differentiation tend to recur among all species comparisons (Supplemental Figure S2). This may indicate that similar regions are under divergent selection among all three species pairs. However, these high *F*_ST_ regions also tend to show valleys in π_B_ and π_W_. This suggests that regions of high *F*_ST_ in our data do not reflect a simple divergence-with-gene-flow model, in which case π_B_ is expected to be higher than in the surrounding homogenized genome (Cruickshank & Hahn, 2014). Neither is it the expected pattern from a model of selection in allopatry, in which π_B_ in high *F*_ST_ regions is similar to π_B_ in low *F*_ST_ regions (Cruickshank & Hahn, 2014; Irwin, 2018; Irwin et al., 2016; T. L. Turner et al., 2005). Therefore, it may be indicative that the current genomic landscape, and regions of high *F*_ST_ in particular, are shaped by recurrent selection (π_B_ lower than in low *F*_ST_ regions; Cruickshank & Hahn, 2014), likely via between-populations selective sweeps followed by differentiation between populations (π_B_ much lower than in low *F*_ST_ regions) (Irwin et al., 2016, 2018).

The geographic landscape was also shaped by recurrent forces at the time sapsuckers began diverging. Pleistocene era glacial fluctuations affected the entire planet, but habitat shifts were especially marked in high latitude regions including northern North America, where kilometers-high ice sheets advanced and receded over the millennia. Sapsuckers, along with other species in this region, were repeatedly isolated into separate refugia with distinct selective pressures before coming back into contact in interglacial periods (L. Natola & Burg, 2018). We suspect that the signatures of recurrent selection or selective sweeps we observe in our sapsucker genome scans are related, whether directly or indirectly, to the recurrent geographic and glacial cycles of the Pleistocene era.

### Z chromosome differentiation

The most striking pattern in our data is the high differentiation along the Z chromosome scaffolds. These patterns fit with well described patterns of accumulation of differentiation on the Z chromosome in birds and other ZW systems, and the X chromosome in XY sex chromosome species (Dean et al., 2015; Henderson & Brelsford, 2020; Irwin, 2018; Qvarnström & Bailey, 2009). The large Z effect, in which the Z chromosome is disproportionately involved in causing inviability or sterility in hybrids (Storchová et al. 2010; reviewed by Irwin 2018), and the fast Z effect describing higher rates of functional change on the Z (Ellegren, 2009; Mank et al., 2010), are particularly well supported. Lower recombination rates and effective population sizes paired with stronger sexual selective pressures, greater drift, and higher mutation rates are known to drive evolution of loci on the sex chromosomes (Axelsson et al., 2004; Charlesworth, 2001; Dean et al., 2015; Irwin, 2018; Mank et al., 2010; Presgraves, 2018). However, considering the three sapsucker species as a whole, the Z chromosome shows less variation than the autosomes do: 0.8% of all autosomal sites are variable, whereas only 0.5% of Z chromosome sites are variable. While the Z chromosome having a smaller effective population size than autosomes can explain some of the reduction in diversity on the Z compared to the autosomes, the ratio of adjusted π_W,Z_ to π_W,A_ is below the theoretical minimum of 0.5625 that can be explained without invoking selection (Charlesworth, 2001; Irwin, 2018). Add to these considerations the finding that π_B_ is also lower on the Z than on autosomes, and we reach the conclusion that the Z chromosome has more recent coalescent times that the autosomes, both within and between species, a pattern likely driven by diversity-reducing selection at multiple timepoints in the past.

The proportion of variant sites themselves are not higher on the Z than on autosomes, but the relative differentiation (*F*_ST_) between haplotypes is far greater on the Z. More than 10% of SNPs on the Z are fixed between red-naped and yellow-bellied sapsuckers, compared to 0.29% among autosomal SNPs. There is precedent for this pattern in the literature. Irwin (2018) compiled a list of 14 bird species comparisons in which the *F*_ST_ values were substantially higher on the Z chromosome than autosomes. There are several hypothesized explanations for this phenomenon. The aforementioned faster rate of functional evolution facilitates the faster accumulation of epistatic incompatibilities between population isolates, causing lower fitness in hybrids which further facilitates differentiation between species at these genomic regions (Irwin, 2018). However, this explanation operates on the premise that SNPs amass on the Z, whereas our data suggest these high *F*_ST_ values with many fixed SNPs are the result of extremely low within population diversity (π_W_) compared to moderately low between-population nucleotide distance (π_B_), and not an accumulation of SNPs. These results provide evidence that coalescence times are more recent in the Z chromosomes than autosomes, despite having higher relative differentiation on the Z (Figure 6).

Postzygotic selection against hybrids can also be invoked with Haldane’s rule, in which deleterious recessives are exposed more readily to selection on the hemizygous Z in females. This is proposed to reduce hybrid fitness, particularly if there are incompatibilities between the Z haplotype of one population and the W or mitochondrial DNA of another (Haldane, 1922; Irwin, 2018; Presgraves, 2002). This accumulation of both differentiation and incompatibilities on the Z chromosome is proposed to play a disproportionate role in the speciation process (reviewed by Irwin 2018), and we might expect patterns of Z differentiation to be most stark on Pleistocene era radiations such as sapsuckers and other incipient boreal species. Because mutations accrue more rapidly and they are more resistant to gene flow on the Z, this would be more apparent in recently diverged species which have experienced recurrent bouts of gene flow. In an earlier analysis of patterns of genetic and phenotypic variation in a three-way hybrid zone of these sapsucker species, Natola et al. (2022) concluded that there is reduced fitness of hybrids, and the present results suggest that incompatibilities on the Z chromosome play a role in that lowered hybrid fitness.

The fixed differences for divergent haploblocks between these sapsucker species might suggest that they are due to chromosomal inversions (or other structural variants), which can result in suppressed recombination and hence accumulation of differentiation between inversion types (da Silva et al., 2019; Huang et al., 2020). We investigated whether the highly differentiated Z chromosomes might be due in part to inversions by inspecting genotype-by-individual plots for two scaffolds covering large regions on the Z (Figure 7). Both of these scaffolds show much evidence for recombination in hybrid birds. Considering this finding together with earlier conclusions that the narrowness of hybrid zones and lack of buildup of hybrids is due to low fitness of hybrids (Natola et al. 2022, 2023), we conclude that the high differentiation across these Z chromosome scaffolds is primarily due to selection against recombinants, rather than nonrecombining inversions.

Hence the strong Z chromosome differentiation between the three sapsucker species may have arisen from a faster functional evolution rate on the Z chromosome (Ellegren, 2009; Mank et al., 2010) combined with epistatic interactions. Substitutions which are neutral, slightly deleterious, or even beneficial in one genome may be strongly deleterious on the genomic or ecological background of another population (Hoekstra et al., 2006; Steiner et al., 2007). These epistatic interactions would be characterized as Bateson-Dobzhansky-Muller Incompatibilities (BDMIs), in which neutral or beneficial evolutionary changes at different loci in separate populations are incompatible upon secondary contact (Bateson, 1909; Dobzhansky, 1936; Muller, 1940). BDMIs, in reducing recombination between haplotypes, also tend to accumulate more epistatic loci, increasing incompatibility in a “snowball effect” (Orr, 1995). Moreover, these may be exacerbated by the disproportionate accumulation of sexually antagonistic sites on the Z chromosome, because selective sweeps in expression modifiers or preference alleles on the Z can accelerate differentiation in sexual traits and preferences between populations, therefore increasing behavioral incompatibilities and further deepening the divide between the species (Irwin, 2018; Pryke, 2010). Indeed, we see evidence that differentiation in sexual traits is associated with loci on the Z chromosome (Natola, 2022).

These patterns of differentiation among sapsucker Z chromosomes could be exacerbated if the various Z haplogroups were each under intense selection unique to a species group. We do see evidence for this scenario in our data. Irwin et al. (2018) attributed patterns of high *F*_ST_ paired with low π_B_ and exceptionally low π_W_ to positive selection causing a selective sweep followed by selective differentiation among populations. In our data, we show π_B_ is significantly lower on Z chromosomes than autosomes, particularly in regions with high *F*_ST_. Within species, π_W_ is significantly lower on the Z chromosome, indicating selective sweeps throughout each Z haploblock, and our π_W,Z_*/π_W,A_ values are lower than the theoretical threshold that could be explained by neutrality, signifying selection must reduce π_W,Z_* in sapsuckers (Charlesworth, 2001; Irwin et al., 2016, 2018). Additionally, sustained homozygosity on haploblocks such as we see within each species on the Z are evidence of strong positive selection on this region (Wooldridge et al., 2022). We propose these large haplotype blocks are driven by strong, divergent selective forces acting upon the different Z haploblocks, with epistasis maintaining the blocks in sympatry via post-zygotic selection against hybrid forms. While we know of few examples that have examined this question directly, we expect that the patterns we observed in sapsuckers could result in large part from a combination of well-characterized, predictable patterns of Z chromosome differentiation and expansive geographical and environmental fluctuations during the Pleistocene era. We expect this mechanism of speciation driven mainly by recurrent divergent selection on Z haplotypes is not unique to sapsuckers, but rather is likely common in incipient high latitude species with chromosomal sex determination.

## Conclusions

During sapsucker differentiation, recurrent selection is implicated by an overall pattern in which regions with high relative differentiation have low pairwise sequence divergence. This pattern is particularly strong on the Z chromosome: relative differentiation is far higher on the Z chromosome than the autosomes, but nucleotide distance is lower on the Z than the autosomes. Recombination in hybrids casts doubt on inversions or other structural variants being the cause of the high Z chromosome divergence. Rather, this pattern might simply result from strong selection on the Z to different species optima combined with epistasis in sympatry causing low fitness of mixed genotypes at the Z chromosome. Relative differentiation between sapsucker species along much of the rest of the genome is low, particularly between red-breasted and red-naped sapsuckers. We suggest that sapsucker species were formed and are maintained in large part due to this differentiation along the Z, which was likely driven by recurrent divergent selection in Pleistocene era glacial refugia.

## Acknowledgements

We thank Sampath Seneviratne and Jocelyn Hudon for sharing blood and tissue samples from hybrid sapsuckers. Thanks to Carla Cicero (Museum of Vertebrate Zoology), Donald McAlpine (New Brunswick Museum), Ray Poulin (Royal Saskatchewan Museum), Garth Spellman (Denver Museum of Natural Sciences), Paul Sweet (American Museum of Natural History), Ildiko Szabo (Beaty Biodiversity Museum), Kevin Winker (University of Alaska Museum) for museum tissues. Thank you to Julia Kreiner and Tom Booker for bioinformatic assistance, and to Kathy Martin, Loren Rieseberg, and Dolph Schluter for feedback on drafts. This research was supported by grants from the Natural Sciences and Engineering Research Council of Canada (Discovery Grants RGPIN-2017-03919, RGPAS-2017-507830, RGPIN-2023-04300), the American Ornithological Society Hesse Award, and the Linnean Society LinnéSys Grant. All protocols were approved by the University of British Columbia Animal Care Committee.

## Author Contributions

LN conceived the project with input from DI. LN sourced samples from museums, extracted DNA, and submitted samples for library preparation. LN prepared raw sequences and ran most analyses. DI converted R scripts to Julia. LN wrote initial drafts with oversight and revisions from DI.

## Data Accessibility

All raw reads will be accessioned on SRA pending publication. All analysis scripts are available in a GitHub repository and will be accessioned for archiving in dryad upon publication acceptance.

## Supplement

**Supplemental Table S1.** Sample metadata, ID includes Irwin Lab ID code, Museum/Band ID includes either museum accession code or CWS bird band record. Museum specifies which museum provided the tissue, or if it was trapped as part of Irwin Lab activities. Species use AOU four-letter species codes RBSA (red-breasted sapsucker), with d indicating *daggeti* and r indicating *ruber* subspecies, YBSA (yellow-bellied sapsucker), RNSA (red-naped sapsucker), and WISA (Williamson’s sapsucker) in allopatry; and YBxRB (yellow-bellied x red-breasted), RNxYB (red-naped x yellow-bellied), RBxRN (red-breasted x red-naped), and RNxYBxRB (red-naped x yellow-bellied x red-breasted) in sympatry. Loc refers to the state or province code where the bird was trapped. Date, Lat, and Long indicate the date and location of sample capture. Sex indicates sex ID as determined by genotypes.

| ID | Museum/Band ID | Museum | Species Zone | Loc | Date | Lat | Long | Sex |
| --- | --- | --- | --- | --- | --- | --- | --- | --- |
| EA22MVZ01 | MVZ:Bird:181742 | MVZ | RBSAd | CA | 2005-01-22 | 38.0807 | -122.1911 | F |
| GG13MVZ01 | MVZ:Bird:191404 | MVZ | RBSAd | CA | 2007-07-13 | 40.4126 | -121.5308 | F |
| GG13MVZ02 | MVZ:Bird:191399 | MVZ | RBSAd | CA | 2007-07-13 | 40.4126 | -121.5308 | F |
| GG13MVZ03 | MVZ:Bird:191401 | MVZ | RBSAd | CA | 2007-07-13 | 40.4126 | -121.5308 | M |
| HF14AMN01 | PRS 3051 | AMNH | YBSA | NY | 2008-06-14 | 42.0413 | -74.8075 | F |
| HG15MVZ01 | MVZ:Bird:182366 | MVZ | RBSAd | CA | 2006-07-15 | 40.3788 | -121.5660 | M |
| IE10S04 | 2251-49108 | Irwin Lab | YBxRB | BC | 2009-05-10 | 55.6969 | -120.2700 | F |
| IE14S02 | 2251-49115 | Irwin Lab | YBxRB | BC | 2009-05-14 | 55.7217 | -121.5736 | M |
| IE15S03 | 2251-49119 | Irwin Lab | YBxRB | BC | 2009-05-15 | 55.7503 | -121.5214 | M |
| IE18S01 | 2251-49122 | Irwin Lab | YBxRB | BC | 2009-05-18 | 55.8267 | -121.7233 | M |
| IE21S02 | 2251-49132 | Irwin Lab | YBxRB | BC | 2009-05-21 | 55.7117 | -121.4925 | F |
| IE24S01 | 2251-49134 | Irwin Lab | YBxRB | BC | 2009-05-24 | 55.4953 | -122.8206 | M |
| IE28S03 | 2251-49147 | Irwin Lab | YBxRB | BC | 2009-05-28 | 55.1958 | -122.6292 | M |
| IF02S01 | 2251-49149 | Irwin Lab | YBxRB | BC | 2009-06-02 | 55.7008 | -122.6267 | M |
| IF03S02 | 2251-49151 | Irwin Lab | YBxRB | BC | 2009-06-03 | 55.2686 | -122.6028 | M |
| IF03S04 | 2251-49152 | Irwin Lab | YBxRB | BC | 2009-06-03 | 55.4611 | -122.6400 | F |
| IF06S02 | 2251-49155 | Irwin Lab | YBxRB | BC | 2009-06-06 | 54.8333 | -123.3836 | M |
| IF13S02 | 2251-49171 | Irwin Lab | YBxRB | BC | 2013-06-09 | 55.0061 | -123.8108 | F |
| IF16S05 | 1841-90618 | Irwin Lab | YBxRB | BC | 2009-06-16 | 54.0481 | -125.3497 | M |
| IF26RABM01 | Z09.8.5 | RABM | RNxYB | AB | 2009-06-26 | 51.8912 | -115.0333 | M |
| JD28BBM01 | TB001883 | BBM | RBSAr | BC | 2010-04-28 | 50.3068 | -122.7821 | M |
| JF14S01 | 1841-90626 | Irwin Lab | RBxRN | BC | 2010-06-14 | 52.1469 | -121.4581 | M |
| JF25RABM01 | Z10.3.9 | RABM | RNxYB | AB | 2010-06-25 | 51.8722 | -115.1365 | M |
| JF26RABM01 | Z10.3.13 | RABM | RNxYB | AB | 2010-06-26 | 51.8059 | -115.2022 | M |
| JF27NBM01 | NBM- 013432 | NBM | YBSA | NB | 2010-06-27 | 45.6878 | -66.1117 | F |
| JF30RABM01 | Z10.13.14 | RABM | RNxYB | AB | 2010-06-30 | 51.8202 | -115.1865 | F |
| JH01NBM01 | NBM- 012066 | NBM | YBSA | NB | 2010-08-01 | 45.6834 | -65.5333 | M |
| KD12DMNS01 | DMNS:Bird:45554 | DMNS | RNSA | SD | 2011-04-12 | 44.4833 | -103.9333 | M |
| KD12DMNS02 | DMNS:Bird:45556 | DMNS | RNSA | SD | 2011-04-12 | 44.4833 | -103.9333 | M |
| KE10DMNS01 | DMNS:Bird:45539 | DMNS | RNSA | SD | 2011-05-10 | 43.6821 | -103.5363 | M |
| KE10DMNS02 | DMNS:Bird:45540 | DMNS | RNSA | SD | 2011-05-10 | 43.6821 | -103.5363 | F |
| KE10DMNS03 | DMNS:Bird:45541 | DMNS | RNSA | SD | 2011-05-10 | 43.6821 | -103.5363 | M |
| KE18S02 | 2251-49198 | Irwin Lab | RBxRN | BC | 2011-05-18 | 52.2206 | -123.0758 | M |
| KE20DMNS01 | DMNS:Bird:34830 | DMNS | RNSA | CO | 2011-05-20 | 40.1146 | -106.0476 | M |
| KE20S02 | 1841-90646 | Irwin Lab | RBxRN | BC | 2011-05-20 | 52.8581 | -124.4383 | F |
| KE22S01 | 1841-90649 | Irwin Lab | RBxRN | BC | 2011-05-22 | 52.5933 | -123.9911 | M |
| KE24S01 | 1841-90653 | Irwin Lab | YBxRB | BC | 2011-05-24 | 52.5331 | -125.4647 | M |
| KE25S03 | 1841-90659 | Irwin Lab | RBxRN | BC | 2011-05-25 | 52.4903 | -125.0964 | M |
| KE27S02 | 1841-90662 | Irwin Lab | YBxRB | BC | 2011-05-27 | 52.6217 | -125.1247 | M |
| KF04S01 | 2251-49187 | Irwin Lab | RBxRN | BC | 2011-06-04 | 51.7361 | -120.8856 | M |
| KF07S01 | 2251-49310 | Irwin Lab | RBxRN | BC | 2011-06-07 | 50.3497 | -115.0022 | M |
| KF07S03 | 2251-49312 | Irwin Lab | RBxRN | BC | 2011-06-07 | 50.2753 | -114.8914 | F |
| KF10S01 | 2251-49317 | Irwin Lab | RBxRN | BC | 2011-06-10 | 49.6467 | -116.4442 | M |
| KF11S02 | 2251-49320 | Irwin Lab | RBxRN | BC | 2011-06-11 | 50.6364 | -118.5450 | F |
| KF13S01 | 2251-49327 | Irwin Lab | RBxRN | BC | 2011-06-13 | 52.7661 | -124.3747 | M |
| KF14S03 | 1841-90675 | Irwin Lab | RBxRN | BC | 2011-06-14 | 52.7853 | -124.2825 | M |
| KF19SO5 | 2251-49346 | Irwin Lab | RBxRN | BC | 2011-06-19 | 52.6317 | -124.6400 | M |
| KF20S07 | 2251-49353 | Irwin Lab | RBxRN | BC | 2011-06-20 | 52.6594 | -124.6256 | M |
| KF21S03 | 1841-90676 | Irwin Lab | RBxRN | BC | 2011-06-21 | 52.5617 | -124.7444 | M |
| KF24RABM01 | Z11.4.30 | RABM | RNxYB | AB | 2011-06-24 | 53.4005 | -117.8685 | M |
| KG01DMNS01 | DMNS:Bird:45552 | DMNS | RNSA | SD | 2011-07-01 | 44.4833 | -103.9333 | F |
| KG07RABM01 | Z11.5.10 | RABM | RNxYB | AB | 2011-07-07 | 51.7774 | -115.2274 | F |
| LE29RABM01 | Z12.2.20 | RABM | RNxYB | AB | 2012-05-29 | 49.7995 | -114.4678 | F |
| LE29RABM02 | Z12.2.7 | RABM | RNxYB | AB | 2012-05-29 | 49.8215 | -114.4246 | F |
| LE30RABM01 | Z12.2.27 | RABM | RNxYB | AB | 2012-05-30 | 50.0243 | -114.0447 | F |
| LF22RABM01 | Z12.3.9 | RABM | RNxYB | AB | 2012-06-22 | 50.6437 | -114.4661 | M |
| MG13RABM01 | Z13.5.7 | RABM | YBSA | AB | 2013-07-06 | 59.9779 | -111.8291 | F |
| MG21DMNS01 | DMNS:Bird:52692 | DMNS | RNSA | CO | 2011-05-21 | 39.9259 | -105.3978 | M |
| ND10L01 | TB001343 | BBM | RBSAr | BC | 2014-04-10 | 49.2422 | -122.9569 | F |
| ND16DMNS01 | DMNS:Bird:43558 | DMNS | RNSA | CO | 2014-04-16 | 39.9337 | -105.3024 | F |
| QD16RABM01 | Z17.10.38 | RABM | YBSA | ON | 2016-04-16 | 45.3443 | -75.9244 | M |
| QD21RABM01 | Z17.10.37 | RABM | YBSA | ON | 2016-04-21 | 45.4180 | -75.7026 | F |
| QF04DMNS01 | DMNS:Bird:47794 | DMNS | RNSA | WY | 2016-06-04 | 41.2333 | -106.7745 | M |
| QF16DMNS01 | DMNS:Bird:46075 | DMNS | WISA | WY | 2016-06-16 | 42.4241 | -105.3365 | M |
| RC06BBM01 | TB001886 | BBM | RBSAr | BC | 2017-03-10 | 49.2236 | -123.1080 | F |
| RC12BBM01 | TB001887 | BBM | RBSAr | BC | 2017-03-12 | 49.2930 | -123.1403 | M |
| RE03NBM01 | NBM- 013265 | NBM | YBSA | NB | 2017-05-03 | 45.2419 | -66.0980 | F |
| RF19DMNS01 | DMNS:Bird:47932 | DMNS | WISA | OR | 2017-06-19 | 44.5388 | -120.6212 | F |
| SE02RSKM01 | RSKM_BIRD_A-9388 | RSKM | YBSA | SK |  | 50.7554 | -104.7088 | M |
| TD02BBM01 | TB001884 | BBM | RBSAr | BC | 2019-04-02 | 49.9572 | -123.1202 | M |
| TF15L03 | 1951-58115 | Irwin Lab | YBSA | BC | 2019-06-15 | 54.1097 | -122.3853 | F |
| TF21L01 | 1951-58120 | Irwin Lab | RN×YB | BC | 2019-06-21 | 52.8331 | -122.6220 | M |
| TF21L02 | 1951-58121 | Irwin Lab | RN×YB×RB | BC | 2019-06-21 | 52.8331 | -122.6220 | F |
| tG15RABM01 | Z94.13.11 | RABM | RN×YB | AB | 1994-07-15 | 51.8300 | -115.2200 | M |
| uE23RABM01 | Z95.9.1 | RABM | RN×YB | AB | 1995-05-23 | 52.2000 | -115.1000 | M |
| uG13RABM01 | Z95.12.4 | RABM | RN×YB | AB | 1995-07-13 | 51.7700 | -115.2200 | M |
| vF04RABM01 | Z96.18.9 | RABM | RN×YB | AB | 1996-06-04 | 49.8200 | -113.9500 | M |
| YE19UAM01 | UAM:Bird:28585 | UAM | YBSA | AK | 2011-07-10 | 64.8570 | -147.8043 | F |

**Supplemental Figure S1.**
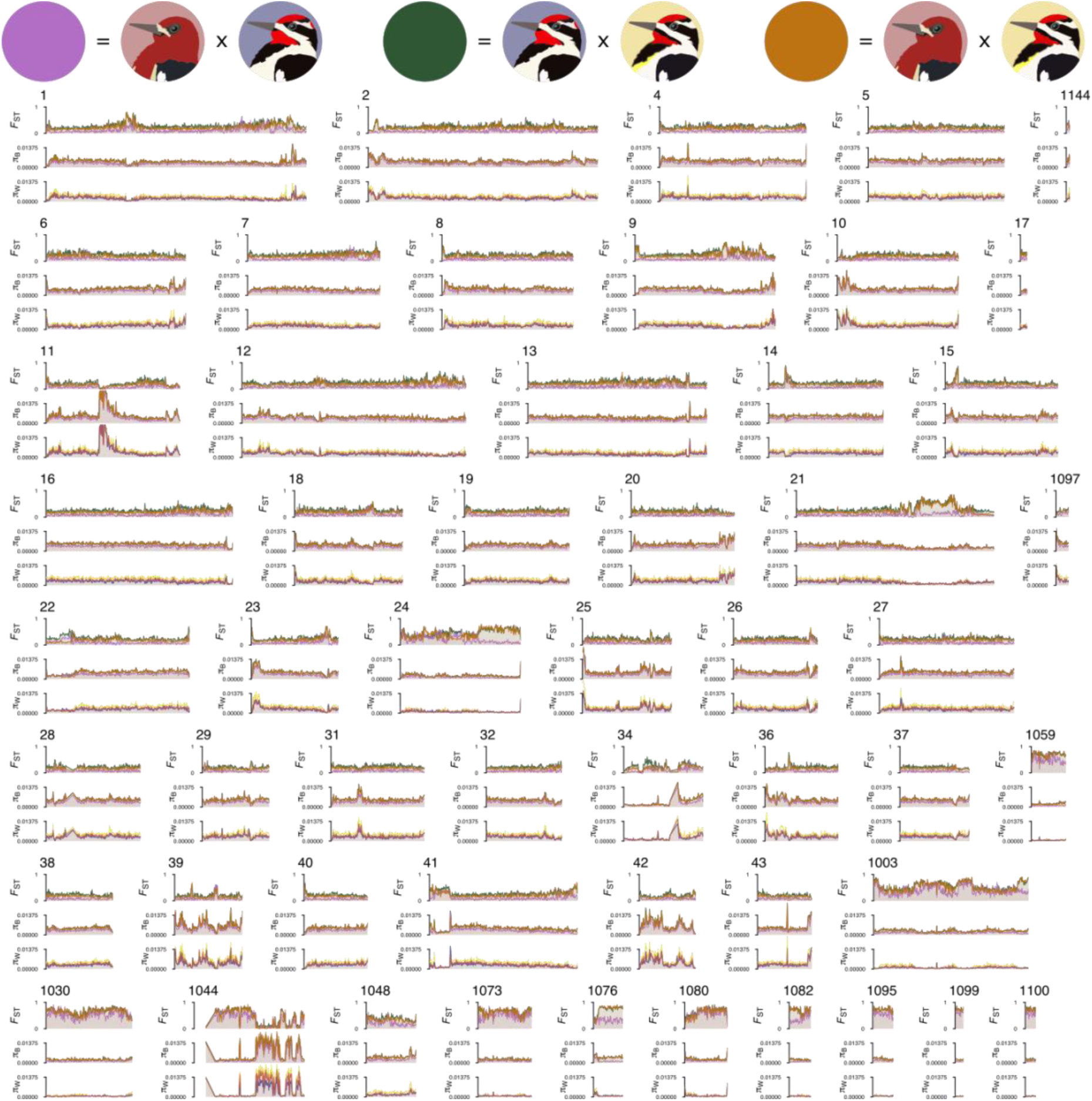
Genomic variation in nucleotide differentiation among sapsuckers for the 45 largest scaffolds and all Z chromosome scaffolds (scaffold IDs > 1000). For each scaffold, graphs show the mean measurement across 50 kb windows for *F*_ST_ (top), π_B_ (middle), and π_W_ (bottom), for all three species/species pairs. The *F*_ST_ and π_B_ species pair lines show red-breasted × red-naped in purple, red-naped × yellow-bellied in green, and red-breasted × yellow-bellied in orange. For π_W_ within species rows indicate red-breasted in red, red-naped in blue, and yellow-bellied in yellow.

**Supplemental Figure S2.**
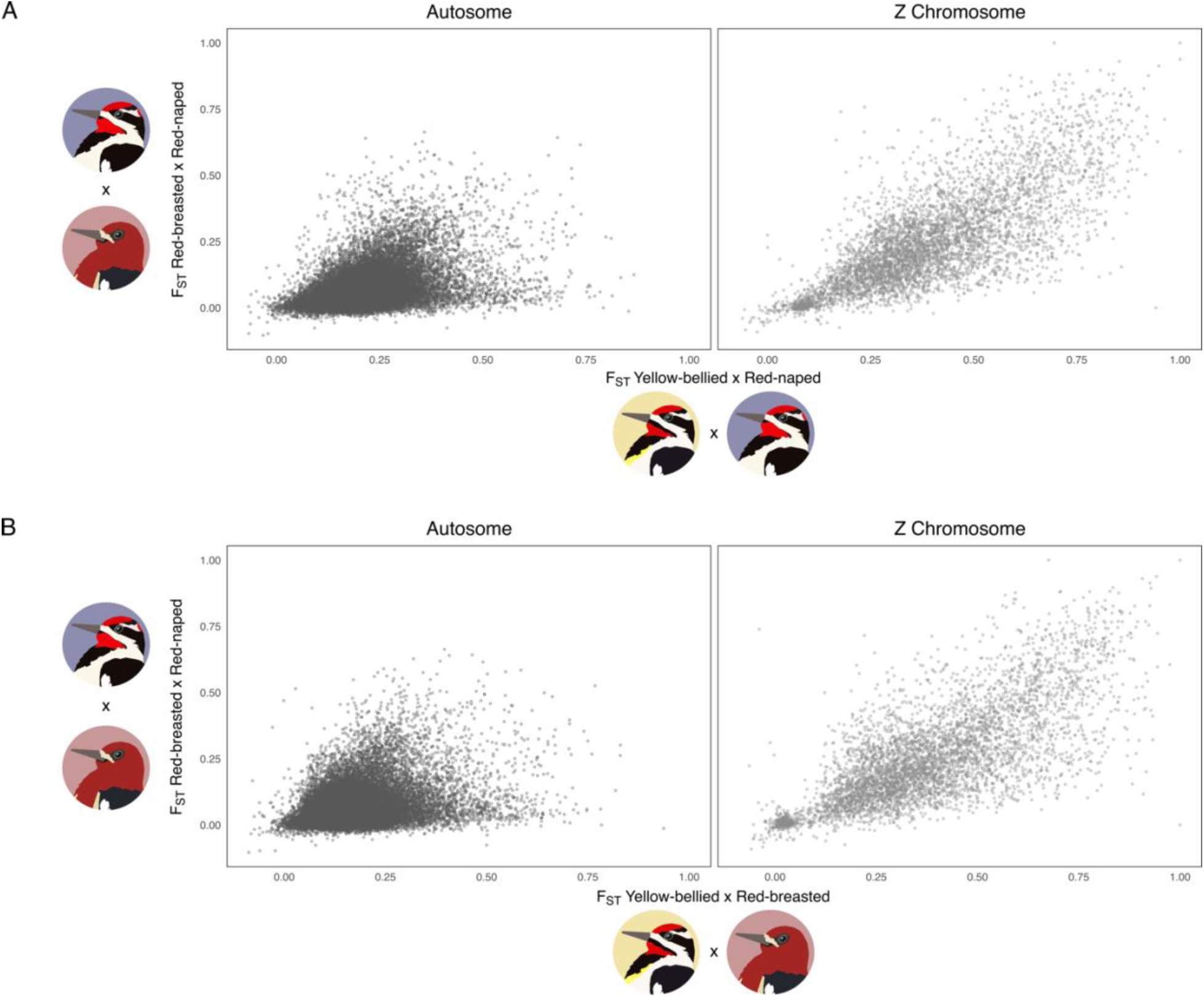
Scatterplots illustrating the relationship between the *F*_ST_ of more distantly related sapsuckers (x-axis) and the *F*_ST_ of the sister species (y-axis) for each 25kb window, paneled separately for autosomal scaffolds (left panels) and Z chromosome scaffolds (right panels). The top panels (A) show how differentiation in Red-breasted and Red-naped sapsuckers compares to differentiation among Yellow-bellied and Red-naped in the same region. The bottom panels (B) show comparisons between Red-breasted and Red-naped *F*_ST_ versus Yellow-bellied and Red-breasted *F*_ST_.

## Notes

### Competing Interest Statement

The authors have declared no competing interest.

### Summary of Updates

This revision includes new explanatory figures, minor additions to statistical methods and figures, and slight reframing of the discussion.

